# Intention to learn modulates the impact of reward and punishment on sequence learning

**DOI:** 10.1101/759639

**Authors:** Adam Steel, Chris I. Baker, Charlotte J. Stagg

## Abstract

In real-world settings, learning is often characterised as intentional: learners are aware of the goal during the learning process, and the goal of learning is readily dissociable from the awareness of what is learned. Recent evidence has shown that reward and punishment (collectively referred to as valenced feedback) are important factors that influence performance during learning. Presently, however, studies investigating the impact of valenced feedback on skill learning have only considered unintentional learning, and therefore the interaction between intentionality and valenced feedback has not been systematically examined. The present study investigated how reward and punishment impact behavioural performance when participants are instructed to learn in a goal-directed fashion (i.e. intentionally) rather than unintentionally. In Experiment 1, participants performed the serial response time task with reward, punishment, or control feedback and were instructed to ignore the presence of the sequence, i.e., learn unintentionally. Experiment 2 followed the same design, but participants were instructed to intentionally learn the sequence. We found that punishment was significantly beneficial to performance during learning only when participants learned unintentionally, and we observed no effect of punishment when participants learned intentionally. Thus, the impact of feedback on performance may be influenced by goal of the learner.

## Introduction

Rewards and punishments play critical roles in shaping human behaviour during learning ^1-3^. Yet despite the importance of valenced feedback for learning, the effects of reward and punishment on performance during the learning of sequencing skills have largely been unexplored ^4^. In the context of sequence learning, two previous studies both demonstrated that punishment improves performance during learning ^5,6^. However, punishment was not beneficial on a continuous sequencing task that required more fine motor control compared to the serial response time task (SRTT) ^6^, suggesting that generalizing the effect of reward and punishment across tasks may be limited. In addition, our prior work has shown that the benefit of feedback to SRTT learning was more pronounced early in training (i.e. after initial exposure the sequence), suggesting that the benefit to learning may be stronger early in learning.

Even within the context of sequence learning, a number of factors affect how the task is learned. For example, when learning in naturalistic settings, humans often have a conceptual understanding of the behavior they intend to learn. Basketball players, for example, have an explicit goal: they should put the ball through the hoop. This intention (putting in the ball in the hoop) is used to guide their actions and shapes the learning process. Despite its importance, to our knowledge, little work has examined the influence of intention in the context of skill learning and feedback. Notably, the intention (i.e. putting the basketball in the hoop) is separable from the physical processes that accomplish this goal (i.e. the specific motor actions involved in shooting a basketball). Thus, it is possible to learn without intention, and it is also possible that what is learned is not known. When learning occurs in the absence of conscious awareness, it is often referred to as “implicit learning ^7^.” Because these features of learning are separable, in this work we use the terms ‘unintentional’ and ‘intentional’ to refer to the goal of the participant during performance, and we use ‘implicit’ and ‘explicit’ to refer to the participant’s subjective awareness of what was learned.

The serial reaction time task (SRTT) is a good task for testing the influence of intentionality on learning. Using the SRTT participants can be instructed to learn the sequence, and the explicit knowledge of the sequence can be assessed post-hoc through the use of awareness tests ^8-10^. Only two previous studies have used the SRTT to study the influence of reward and punishment on sequence learning. In our previous study, participants were not told about the presence of a sequence and demonstrated implicit acquisition when tested at 30-days post-learning ^6^, and in the study by Wachter and colleagues, participants were excluded if they achieved explicit awareness of the presence of the sequence ^5^.

Given the central importance of both reward and punishment and intention to skill learning, we therefore investigated whether the intention to learn modulates the effect of valenced feedback on SRTT performance. The present study comprises two experiments in which participants performed the SRTT augmented with reward, punishment, or control feedback. In the first experiment (the unintentional learning condition), participants were told that a sequence would be present but to ignore the sequence during training, and to continue press buttons as fast and accurately as possible. In the second experiment (the intentional learning condition), participants were told to do their best to learn the sequence (Figure 1). Sequence and random blocks were interleaved, so that we could continuously evaluate sequence knowledge expression. Based on prior work which showed that both reward and punishment enhanced early learning compared to control feedback, we were particularly interested in the difference between the intentional and unintentional learners during the early learning period.

**Figure 1.**
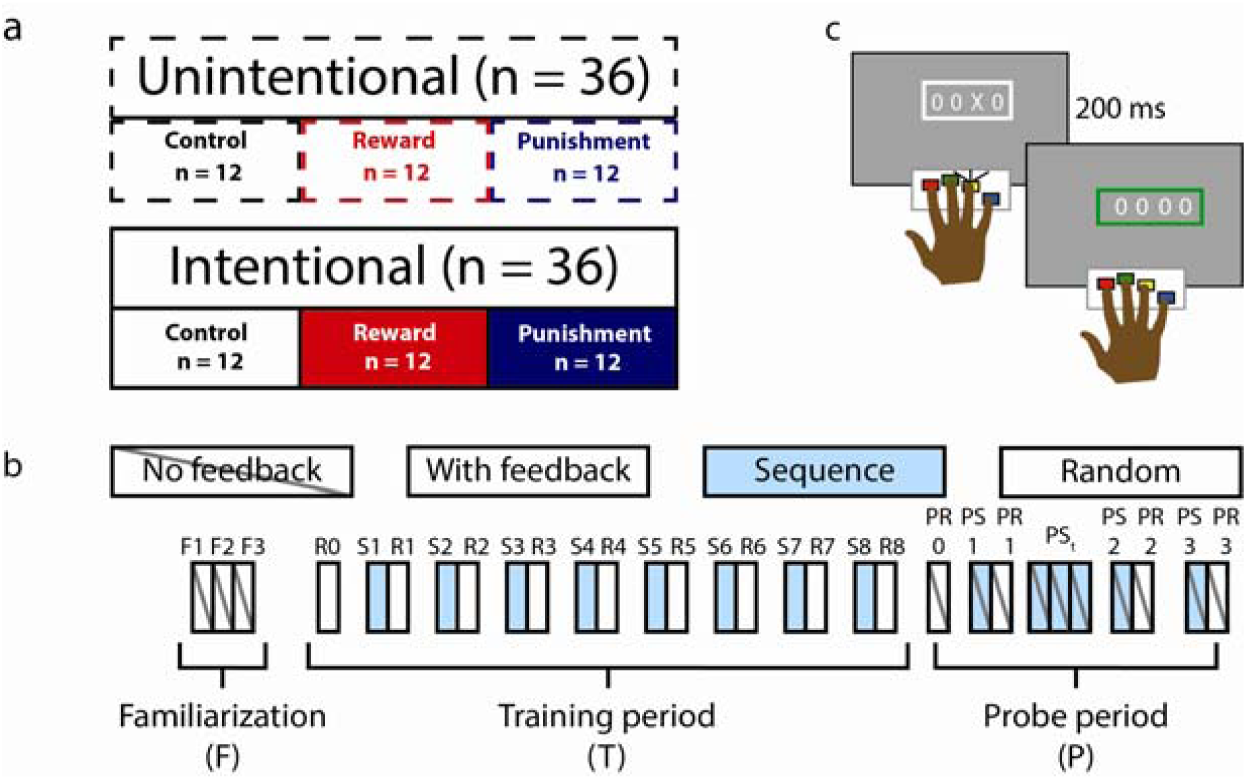
Experimental design. (a) The study consisted of two experiments, which differed based on the instruction given to the participants. Participants performed the SRTT and were told either to learn the sequence intentionally (Experiment 2) or to try to ignore the sequence (Experiment 1). (b) The SRTT took place over 30 blocks. The first three blocks of the task (familiarization blocks [F]) contained no feedback and the stimulus appeared according to a random-sequence. The training period began at the fourth block (random-sequence [R]), at which point feedback was introduced. Beginning in the fifth block, the block order altered such that fixed- and random- sequence blocks were presented in an alternating order. Participant’s RTs during in each fixed-sequence [S] block to the subsequent random-sequence block indicated skill knowledge expression. After the 17th block (eighth fixed-/-random doublet), the immediate retention probe began. During the immediate retention probe, participants performed a random-sequence block without feedback. They then performed a fixed-sequence block followed by a random block, and comparison between these two blocks indicated retention. Participants then performed four consecutive fixed-sequence blocks, followed by a random block. Finally, participants performed a fixed- and fixed-random block. c. In both experiments, participants performed the SRTT with feedback, which was contingent on their speed and accuracy.

## Results

The present study consisted of two experiments that examined intentional and unintentional learning, respectively. The results for each experiment are first presented separately and then the two experiments are formally compared in the final section.

### Experiment 1 – Unintentional learning

#### Punishment improves sequence knowledge expression during unintentional learning

The reward, control, and punishment groups did not differ in their RT during the familiarization period (Blocks F1-F3; One-way ANOVA: F_(2,33)_ = 1.372, p = 0.27). We first sought to determine whether training with reward or punishment influenced the acquisition and expression of sequence knowledge (indexed by comparing RTs during fixed- and random-sequence blocks) during the training period. To this end, we compared the baselined RT (i.e., improvement relative to the three familiarization blocks to control for individual differences in RT) during the training period (S1 through R8) using a repeated measures ANOVA with Feedback type (Reward/Punishment/Control), Sequence type (Fixed-/Random-sequence), and Block (1-8) as factors (Figure 2).

**Figure 2.**
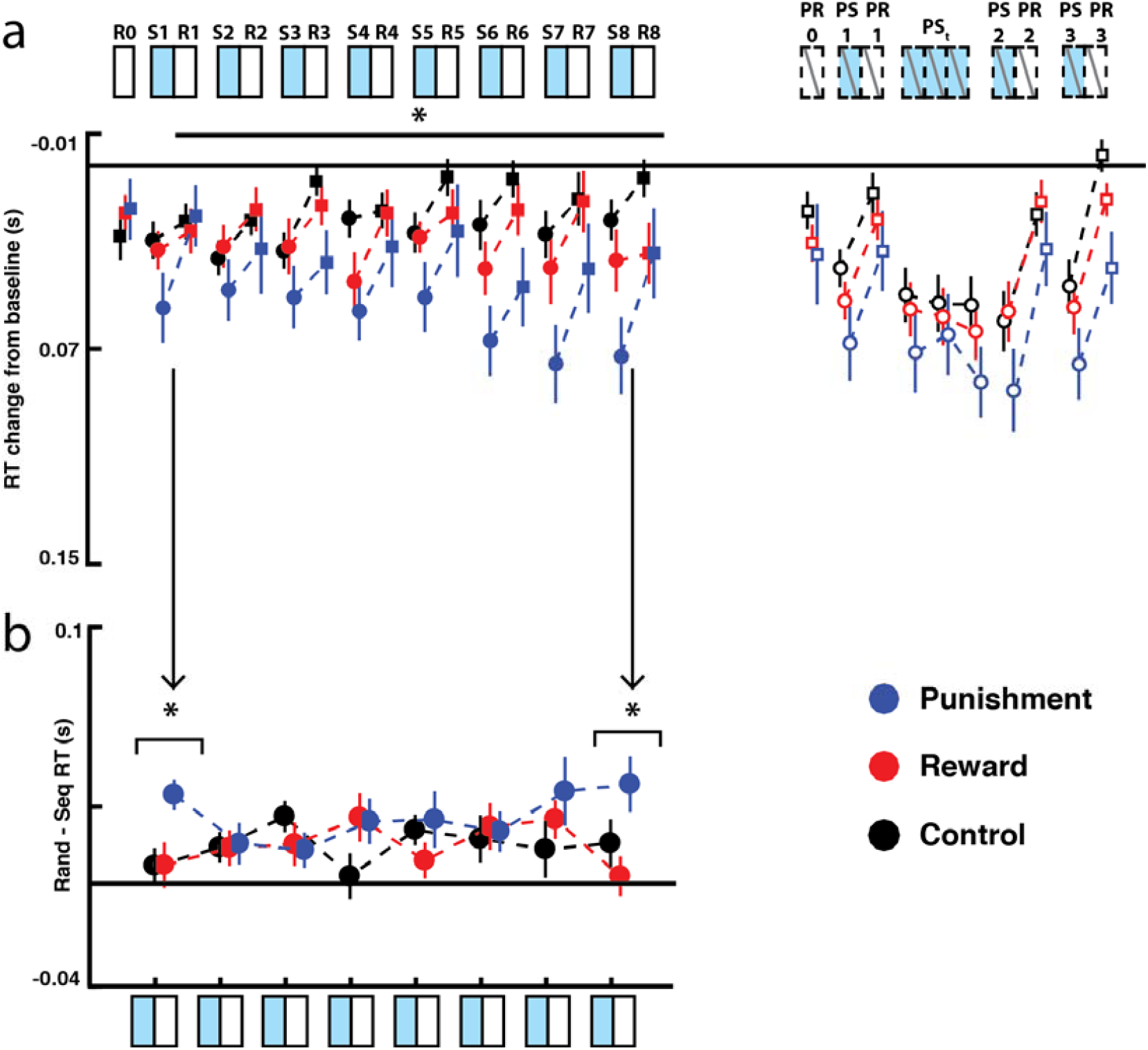
Punishment improves unintentional sequence learning. (a) The participant’s performance during the training and immediate test conditions relative to their performance during the familiarization blocks. (b) Difference between fixed- and random-sequence RTs over the training period. The punishment group evidenced more sequence knowledge during the first block of training with feedback (S1 v R1), as well as during the final block of training (S8 v R8). Asterisks indicate significant difference (p < 0.05). Error bars show SEM. R = training random-sequence, S = training fixed-sequence, PR = probe random-sequence, PS = probe fixed-sequence.

This analysis revealed that the difference in RT between the fixed- and random-sequence blocks varied among the feedback groups over the course of learning (three-way interaction, Feedback type × Sequence type × Block: F_(7,231)_ = 1.741, p = 0.049, η^2^ = 0.01). Follow-up analyses revealed the punishment group evidenced greater sequence knowledge during the first block of training (S1 versus R1) compared to the reward and control groups (Punishment vs Reward: t_(23)_ = 2.77, p_corr_= 0.026; Punishment vs Control: t_(23)_ = 2.79, p_corr_ = 0.026). The punishment group also evidenced greater sequence knowledge during the final block of training (S8 v R8) with feedback compared to the control group but not to the reward group (Punishment versus Control: t_(23)_ = 2.81, p_corr_= 0.022; Punishment versus Reward: t_(23)_ = −1.013, p_corr_= 0.19). There were no significant differences in performance between the feedback groups for any other block. There was no difference between the reward and control groups at the beginning (t_(23)_ = 0.02, p_corr_= 0.98) or end of training (t_(23)_ = 1.0, p_corr_= 0.57), suggesting that punishment, rather than reward, is beneficial to sequence knowledge expression during unintentional sequence learning.

In addition to the difference in performance across the reward, control, and punishment groups that differed by sequence type, the overall RT differed across the feedback groups over time (Feedback type × Block interaction: F _(11,183)_= 2.47, p = 0.006, η^2^ = 0.018). Although we did not have any *a* priori expectation that RT would differ across the blocks, we conducted exploratory post-hoc analyses to better understand these effects. These post-hoc analysis revealed that the participants in the punishment group performed significantly faster than the control group during Block 3 (Punishment v Control; t_(23)_ = 2.47, p_corr_ = 0.046) and Block 7 (Punishment v Control; t_(23)_ = 2.67, p_corr_ = 0.033). The reward group was not significantly faster than the control group during any block (all ps > 0.05).

Across the training period, there was no effect of feedback type on accuracy during the training period with feedback (Main effect of feedback type: F_(2,33)_ = 2.34, p = 0.112; Group × Block: F_(14,231)_ = 0.97, p = 0.478; Group × Sequence type: F_(2,231)_ = 0.58, p = 0.56; Group × Sequence type × Block: F_(14,231)_ = 1.17, p = 0.30). These data are presented in Supplemental table 1.

#### Short-term retention is not impacted by valenced feedback during unintentional learning

We next sought to determine whether valenced feedback impacted short-term sequence-knowledge expression. First, to examine immediate skill performance, we compared RTs during the first fixed- and random-sequence blocks presented in the first probe block (PS1 v PR1) using a repeated measures ANOVA with Feedback type (Reward/Control/Punishment) and Sequence type (Fixed-/Random-sequence) as factors. All groups expressed skill knowledge during the immediate probe (Main effect of Sequence type, F _(1,33)_ = 39.01, p < 0.001, η^2^ = 0.192; Fixed-vs Random-sequence: t_(35)_ = 6.40, p < 0.001). There was no effect of feedback type on skill knowledge expression during the first probe block (Main effect of Feedback: F_(2,33)_ = 1.10, p = 0.36; Feedback x Sequence interaction: F_(2,33)_ = 0.616, p = 0.55).

Punishment has previously been shown to increase relearning rate in visuomotor adaptation tasks, and we sought to determine whether this phenomenon also occurred in sequence learning. To examine this question, we compared the RTs during the four-consecutive fixed-sequence blocks in the probe period (PSt1-3 and PS2) using a repeated measures ANOVA with Feedback type (Reward/Punishment/Control) and Block (PSt1-PS2) as factors. There was no effect of Feedback type (F_(2,33)_ = 1.02, p = 0.37), Block (F_(3,99)_ = 2.098), p = 0.11), or Feedback type x Block interaction (F_(6,99)_ = 0.95, p = 0.46). There was also no difference between the groups during the second probe block (after the retraining probe, PS2 v PR2; Feedback type x Sequence type: F_(2,33)_ = 0.577, p = 0.57). While all groups demonstrated learning (retraining probe – Main effect of Sequence type: F_(1,33)_ = 68.43, p < 0.001, η^2^ = 0.203), there was no difference in skill knowledge across the groups in the final skill knowledge probe (Feedback type × Sequence type: F_(2,33)_ = 0.577, p = 0.57). In the final retention probe (PS3 v PR3), the punishment group was significantly faster than the control group (Main effect of Feedback type (F_(2,33)_ = 3.942, p < 0.029, η^2^ = 0.193); Punishment versus Control: t_(23)_ = 2.762, p_corr_ = 0.028, Punishment versus Reward: t_(23)_ = 1.817, p_corr_ = 0.156, Reward versus Punishment: t_(23)_ = 0.945, p_corr_ = 0.352). However, this was not specific to the trained sequence, as all groups showed learning (Main effect Sequence type F_(1,33)_ = 57.512, p < 0.001, η^2^ = 0.240; fixed-sequence versus random-sequence: t_(35)_ = 7.702, p_corr_ < 0.001), and there was no difference between the feedback groups in terms of sequence knowledge expression (Feedback type × Sequence type F_(2,33)_ = 0.468, p = 0.63).

#### Feedback does not alter awareness during unintentional learning

Lastly, we sought to determine whether the addition of reward or punishment altered conscious awareness during unintentional skill learning. To assess the degree of implicit/explicit knowledge, we utilized two different measures. First, participants indicated knowledge via verbal report. For this test, we asked participants to verbally recall as much of the sequence as they were confident in producing. The number of sequential triplets included was compared across the groups using a one-way ANOVA. The feedback groups did not differ in the number of sequential items reported (F_(2,33)_ = 1.62, p = 0.21).

Second, we used the process dissociation procedure ^10^. To assess any purely implicit bias, before the presence of the sequence was reaffirmed, we asked participants to perform a series of button presses and try to be as random as possible (free generation). Then, participants were asked to include or exclude the sequence during a series of 100 button presses. One participant was removed from the analysis due to data collection failure. We calculated the area under the curve which describing the number of sequential items generated (doubles, triples, quads, etc) for each set of button presses and compared across the groups on this measure using a one-way ANOVA (Figure 3).

**Figure 3.**
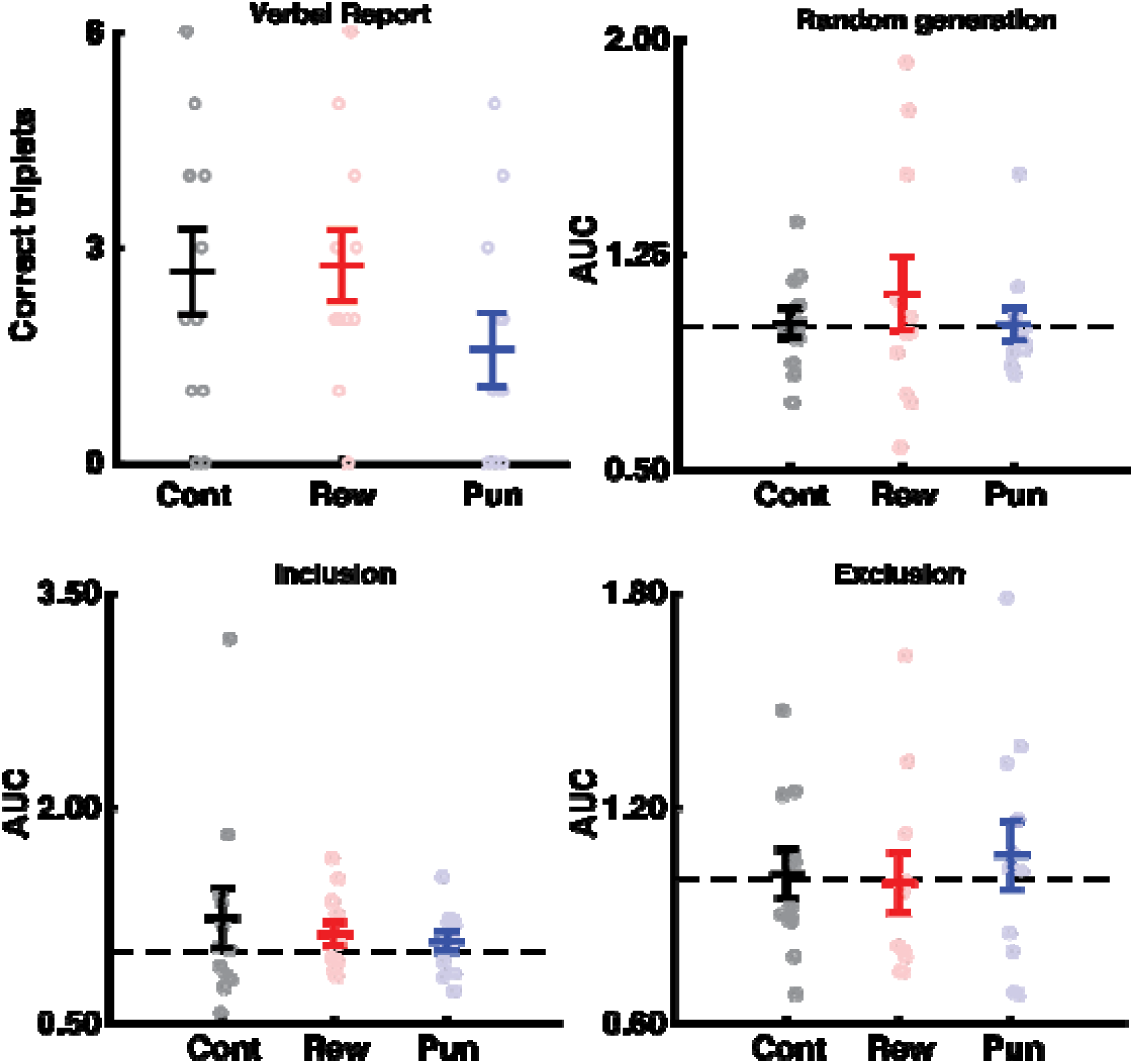
Feedback did not impact awareness when learning unintentionally. There was no difference between the Feedback groups with respect to explicit knowledge formed during the unintentional learning condition based on verbal report, as well as process dissociation procedure inclusion, exclusion, or free generation rates. Error bars show SEM. Colors: Black = control, red = reward, blue = punishment. Dashed lines indicate chance-level as determined by 1000 Monte-Carlo simulations.

Using this procedure, we found no difference in sequence inclusion on any set of presses between the control, reward, and punishment groups (Free generation: F_(2,32)_ = 0.528, p = 0.95; Inclusion: F_(2,32)_ = 0.400, p = 0.68; Exclusion: F_(2,32)_ = 0.233, p = 0.79).

#### Unintentional learning summary

In summary, when learning unintentionally, we found that participants training with punishment demonstrated significantly more skill knowledge during the acquisition period than participants training with reward or control feedback. The feedback given during the training period did not influence participant’s ability to express sequence knowledge immediately after training after feedback was removed and did not systematically modulate the implicitness of the sequence acquisition.

### Experiment 2 – Intentional learning

#### Punishment does not benefit intentional sequence learning

To determine whether punishment similarly benefitted intentional sequence learning, we performed the same experiment described above, but instructed participants to intentionally learn the sequence. Data from this experiment are shown in Figure 4. There was no difference between the feedback types during the Familiarization period (F1-F3; one-way ANOVA: F_(2,33)_ = 1.094, p = 0.34). We next examined the impact of feedback on performance during the training period (S1-R8) using a repeated measures ANOVA with Feedback type (Reward/Punishment/Control), Sequence type (Fixed-/Random-sequence), and Block (1-8) as factors. When instructed to learn intentionally, all feedback types showed sequence knowledge improvement over time (Sequence type × Block interaction: F _(7,231)_ = 4.81, p < 0.001, η^2^ = 0.02; fixed-versus random-sequence difference in Block 1 versus Block 8, t_(35)_ = 3.781, p < 0.001). However, feedback type did not impact overall performance or sequence knowledge (Main effect of Feedback type: F_(2,33)_ = 1.13, p = 0.34; Feedback type × Block: F_(14,231)_ = 0.903, p = 0.56; Feedback type × Sequence type: F_(2,33)_ = 0.244, p = 0.79; Group × Sequence type × Block: F_(7,231)_ = 1.36, p = 0.18; Figure 4). Thus, in contrast to the benefit of punishment found in unintentional learning, when instructed to learn intentionally, the punishment group did not express sequence knowledge earlier than the reward and control groups.

**Figure 4.**
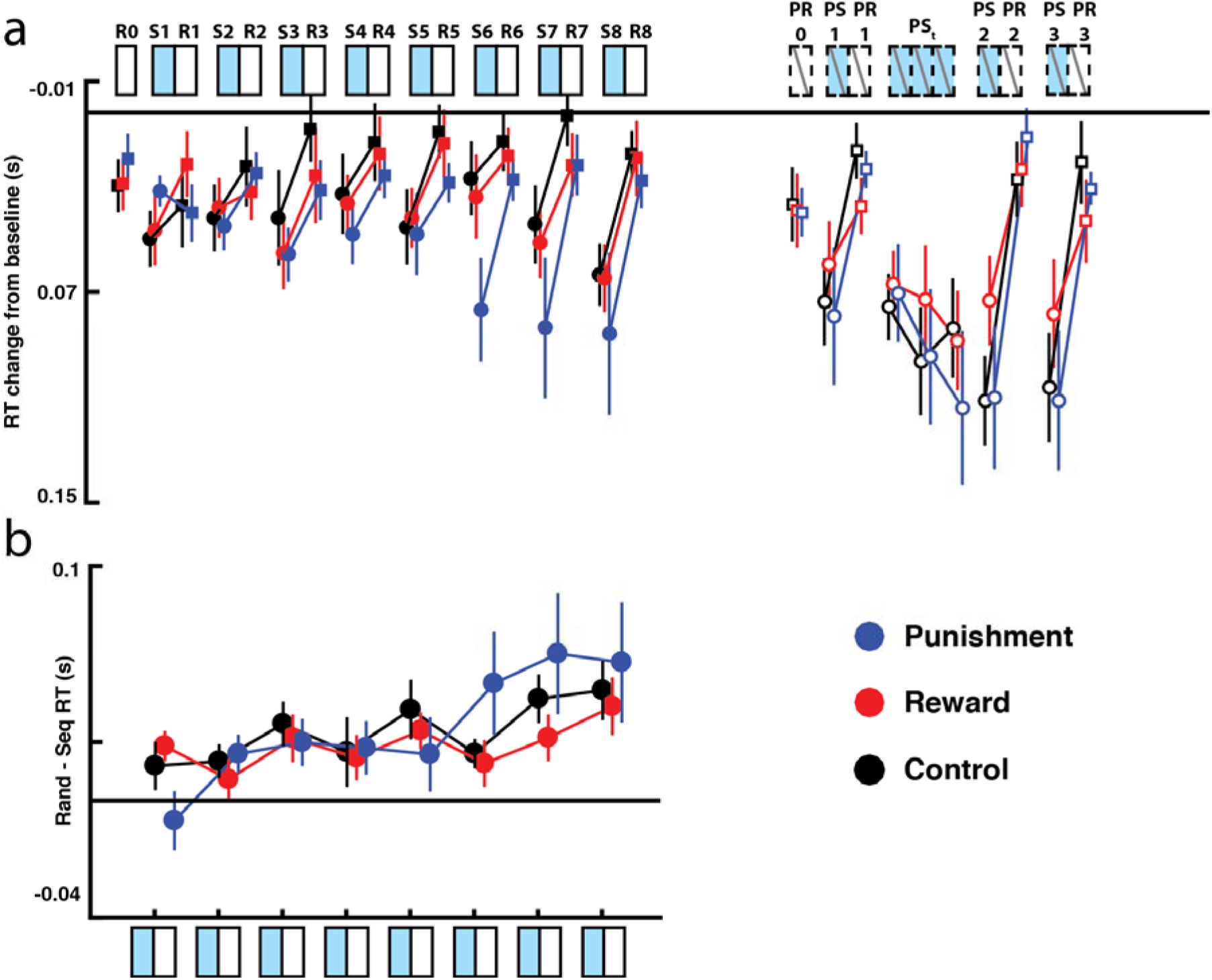
Feedback does not impact performance during intentional sequence learning. (a) All feedback groups showed robust learning when instructed to learn the sequence (b) Difference in RTs during fixed- and random-sequence blocks across the learning period. All groups showed equivalent development of sequence knowledge. Error bars show SEM. R = random-sequence, S = fixed-sequence, PR = probe random-sequence, PS = probe fixed-sequence.

#### Feedback does not alter retention when learning intentionally

We next investigated whether feedback modulated retention or relearning after intentional sequence learning. All groups expressed skill knowledge during the first probe block (PS1 v PR1; Main effect of Sequence type, F_(1,33)_ = 26.00, p < 0.001, η^2^ = 0.209; fixed-vs random-sequence: t_(35)_ = 5.09, p < 0.001). However, there was no effect of feedback type on skill knowledge expression during the first probe block (Main effect of Feedback type: F_(2,33)_ = 1.09, p = 0.35; Feedback type × Sequence type: F_(2,33)_ = 0.26, p = 0.80).

When we examined relearning (PSt 1-3 and PS2), we found that the control, reward, and punishment groups all showed improvement during the relearning period (Main effect of Block: F _(3,99)_ = 9.105, p < 0.001, η^2^ = 0.038). However, there was no effect of feedback on relearning (Main effect of Feedback type: F_(2,33)_ = 0.68, p = 0.51; Feedback type x Block interaction: F_(6,99)_= 0.95, p = 0.46). Similar to the unintentional learning, the control, reward, and punishment groups all showed sequence knowledge after relearning during probe block 2 (PS2 v PR2; Main effect of Sequence type: F_(1,33)_ = 68.475, p < 0.001, η^2^ = 0.365), but there was no impact of Feedback type on the sequence knowledge expressed during probe block 2 (Feedback type × Sequence type: F_(2,33)_= 0.50, p = 0.61). In the final retention probe (PS3 v PR3), all groups showed learning (Main effect Sequence type F _(1,33)_ = 30.97, p < 0.001, η^2^ = 0.312), but there was no effect of feedback on performance or evidenced knowledge (Main effect of Group: F_(2,33)_= 0.79, p = 0.47; Sequence type × Group F_(2,33)_= 1.05, p = 0.36).

#### >Awareness was not modulated by feedback when learning intentionally

Finally, we sought to determine whether the degree of explicit awareness after intentional sequence learning was modulated by feedback (Figure 5). The feedback groups did not differ in the number of sequential items verbally reported (F_(2,33)_ = 0.36, p = 0.70). Using the process dissociation procedure described above, we found no difference in sequence inclusion on any set of presses (Random: F_(2,32)_ = 0.35, p = 0.70; Inclusion: F_(2,32)_ = 0.1.48, p = 0.24; Exclusion: F_(2,32)_ = 0.7, p = 0.50).

**Figure 5.**
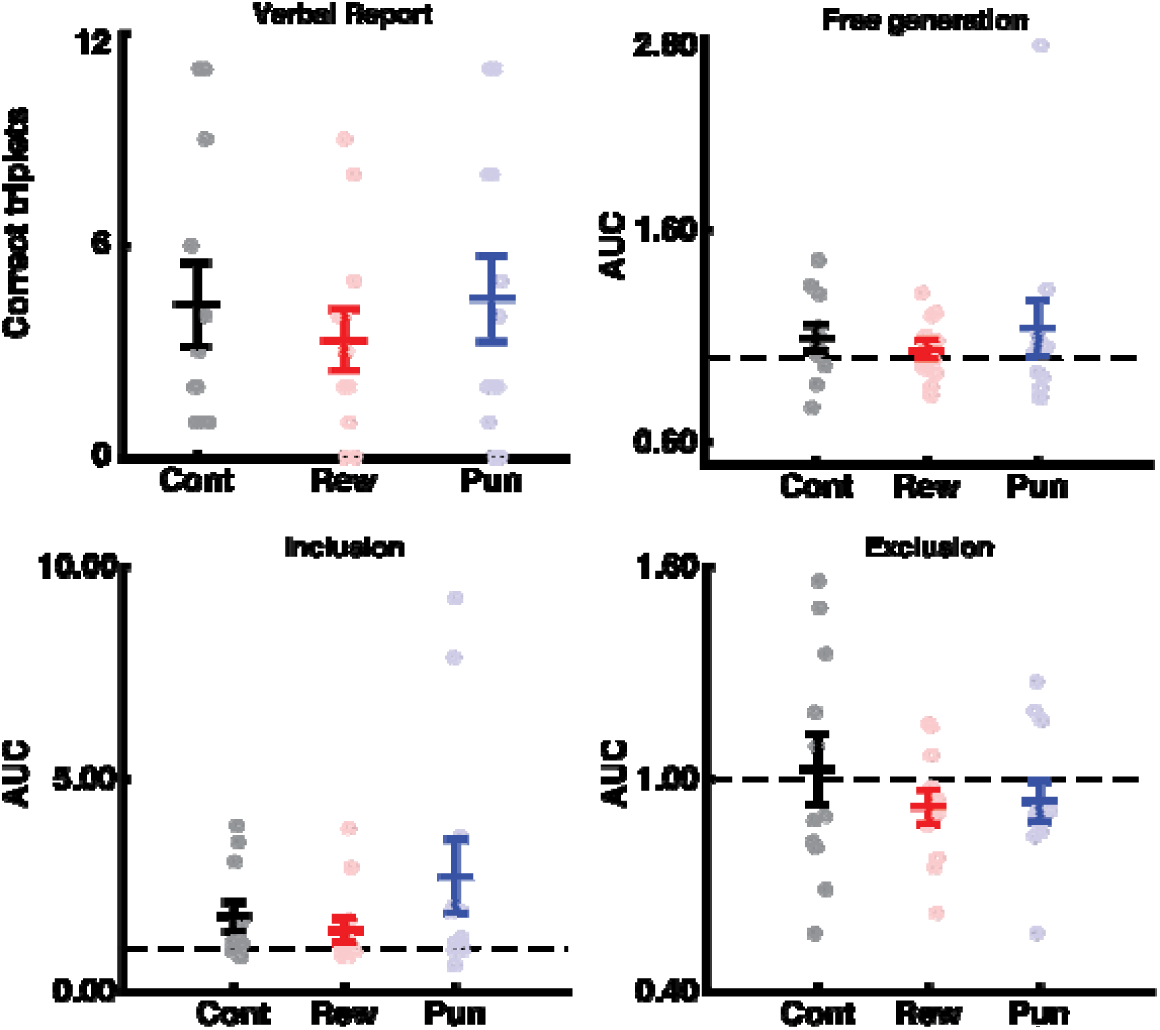
There was no difference between the feedback groups with respect to explicit knowledge formed during intentional sequence learning. Error bars show SEM. Colors: Black = control, red = reward, blue = punishment. Dashed lines indicate chance-level based on 1000 Monte-Carlo simulations.

#### Intentional learning summary

In summary, the type of feedback given during training did not impact performance when participants learned intentionally. The type of feedback also did not impact retention or conscious knowledge expression.

### Punishment specifically benefits unintentional learning

To formally test for a difference between the intentionality conditions, we directly compared the performance during the training period from Experiments 1 and 2 (S1-R8) using a repeated measures ANOVA with Intentionality (Intentional/Unintentional), Feedback type (Reward/Punishment/Control), Sequence type (Fixed-/Random-sequence), and Block (1-8) as factors. The full ANOVA investigating whether intentionality influenced the effect of valenced feedback on sequence learning rate was not significant (4-way interaction, Intentionality × Feedback type × Sequence type × Block; F (28,462) = 1.483, p = 0.055, η^2^ = 0.028). However, based on our a *priori* hypothesis that intentionality would specifically impact early learning, we conducted a repeated measures ANOVA examining Block 1 (S1 v R1). For this ANOVA we considered Intentionality (Intentional/Unintentional), Feedback type (Reward/Punishment/Control), and Sequence type (Fixed-/Random-sequence) as factors. This analysis revealed that intentionality did impact the influence of feedback valence on early knowledge expression (three-way interaction, Intentionality × Feedback type × Sequence type; F_(2,66)_ = 7.21, p = 0.001, η^2^ = 0.117). Follow-up tests revealed that intentionality specifically modulated the influence of punishment during Block 1 (S1 v R1; Figure 6). The punishment group learning unintentionally evidenced more sequence knowledge during Block 1 compared to punishment group learning unintentionally (Unintentional Punishment vs Intentional Punishment: t_(22)_ = 3.241, p = 0.0038). The unintentional punishment group did not perform significantly differently to the intentional reward or intentional control groups (Unintentional Punishment vs Intentional Reward: t_(22)_ = 1.350, p = 0.19; Unintentional Punishment vs Intentional Control: t_(22)_ = 1.733, p = 0.10). Intentionality did not influence sequence knowledge expression during Block 1 in either the reward or the control groups (Unintentional Control vs Intentional Control: t_(22)_ = 0.658, p = 0.51; Unintentional Control vs Intentional Reward: t_(22)_ = 0.583, p = 0.56; Unintentional Reward vs Intentional Reward: t_(22)_ =1.531, p = 0.14; Unintentional Reward vs Intentional Control: t_(22)_ = 0.58, p = 0.566). The feedback groups did not differ at any other blocks. Thus, these data support the conclusion that intentionality specifically modulates the impact of punishment on early learning.

**Figure 6.**
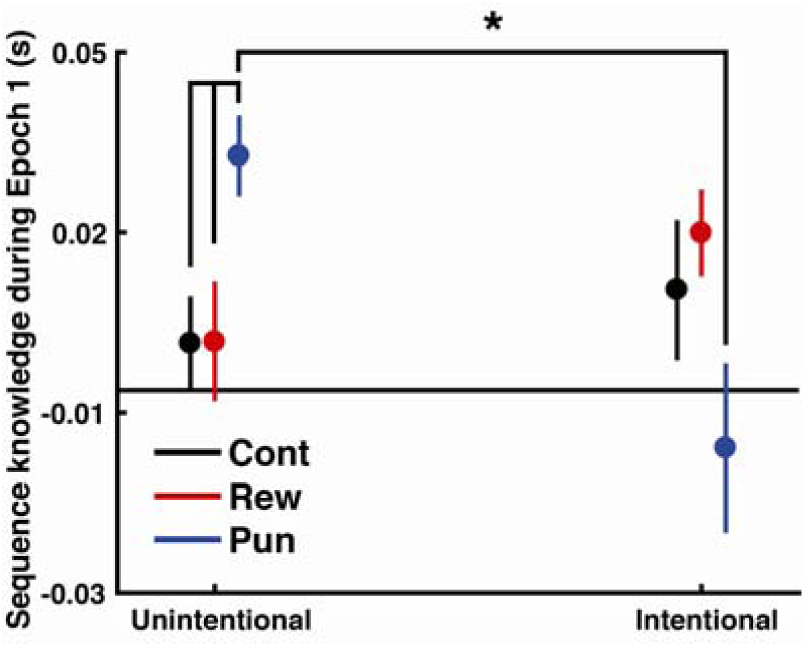
Comparison of sequence knowledge expression (S1 v R1) across the feedback groups in unintentional versus intentional learning. The group of participants learning unintentionally with punishment performed significantly better than the group of participants learning intentionally with punishment, as well as the reward and control groups learning unintentionally. Asterisks indicate a significant difference (p < 0.05). Error bars show SEM.

With respect to awareness measures, intentional learning fostered significantly greater explicit awareness, as evidenced by a greater number of sequence items generated during the verbal report (F_(1,66)_= 6.452, p = 0.013, η^2^ = 0.086; Intentional v Unintentional: t_(68)_= 2.54, p = 0.013) and inclusion rate during the process dissociation procedure (F_(1,66)_= 6.39, p = 0.014, η^2^ = 0.084; Unintentional v Intentional: t_(68)_= 2.50, p = 0.014). There was no main effect or interaction with feedback group in the awareness tests for either experiment (all ps > 0.3).

## Discussion

In this study we investigated whether the intention to learn interacts with the effects of valenced feedback on performance during sequence learning. We found that punishment enhanced early sequence knowledge expression when participants learned unintentionally (Experiment 1). However, when learning intentionally, all groups performed equally well (Experiment 2). Based on our prior work, we expected that punishment would specifically benefit sequence knowledge during early learning ^6^, and direct comparison of the results of the two experiments revealed that punishment was, indeed, beneficial to sequence knowledge formation in the early phase of unintentional sequence learning. These results suggest that intentionality may influence the impact of valenced feedback given during performance.

The effect of reward and punishment on performance during learning has previously been characterized with a simple heuristic: punishment being beneficial to performance, while reward benefits memory retention ^4,5,11^. A previous study by our group demonstrated that this simple heuristic is not valid in all cases. Specifically, we showed that the impact of valenced feedback depends on the task being performed: while punishment benefited performance on the SRTT when learning unintentionally, punishment was detrimental to performance on a different motor learning task (the force tracking task). In the present study, we additionally show that even when participants are performing the same task, altering the intention of the participant may influence the effect of reward and punishment. This result further complicates the use of the simple heuristic outlined above even within the same task. Together, these two results suggest that valenced feedback interacts with skill learning in a complex manner.

### Why might intentionality modulate the impact of feedback?

The central question raised by this study is why punishment benefits unintentional learning but not intentional learning. A critical component of the intentional learning is the added cognitive load introduced by intentionally learning the sequence. The ‘cognitive load’ may be due to several factors, but two examples are i) the added ‘dual-task’ of learning to maximizing the amount earned, and ii) the added performance monitoring during negative feedback that may cause ‘choking under pressure’ in intentional learning tasks, or iii) fatigue caused by intensive training. Because the present task was self-paced and took place over a short duration, the third possibility is unlikely to cause the results observed here. In contrast, either, or both, of the former points might explain the different effects of punishment on intentional and unintentional sequence learning.

#### Introducing a competitive task

It is possible that the goal of learning the sequence acts as a distractor task that competes with the primary task of pressing buttons quickly and accurately. While participants in the intentional and unintentional learning conditions share the goal of maximizing the total money earned during the experiment, participants learning intentionally are also compelled to detect and learn a sequence, while participants learning unintentionally are not. The added cognitive load in the intentional condition, then, may be similar to dual-task learning. In dual-task learning, participants are forced to learn while simultaneously performing another task that competes for attention, for example counting tones ^12-16^. Under dual-task conditions, sequence knowledge may take longer to be expressed ^12-14^, and the detrimental effect of the added cognitive load may be more extreme when participants engage explicit strategies ^16^.

Although it should be noted that visuomotor adaptation is generally considered to be a separate process from sequence learning, complementary evidence comes from the visuomotor adaptation literature (for an in-depth discussion regarding the SRTT and its relationship to other motor-learning tasks, see a recent review from Krakauer and colleagues^17^). Similar to prior work in sequence learning ^5,11^, prior work using motor adaptation tasks have found that reward benefits retention, while punishment benefits performance during training ^18-21^. For example, it has been found that the beneficial effects of reward-feedback to retention in visuomotor adaptation tasks appear to depend on explicit processes, and a distractor-task that disrupts explicit processes completely abolishes learning when participants must use reward-based feedback to adapt ^18,22,23^. In addition, artificially increasing noise during adaptation learning is detrimental to learning through reinforcement, which is also consistent with the importance of explicit processes to adaptation learning ^24^. While studies of visuomotor adaptation have not explored the role of explicit processes when learning with punishment, this could be an interesting avenue for future research.

#### Engaging alternative cognitive strategies

A second possible source of cognitive load that may account for the difference between intentional and unintentional learning is that the engagement of the intentional learning strategy in the context of negative feedback invokes meta-cognitive processes (e.g., performance monitoring) that interfere with learning ^25-28^. One study has examined the role of performance monitoring in the context of the SRTT when learned unintentionally (implicitly). In this study, a control condition (no monitoring) was compared i) to an outcome-pressure condition, where participants were told that performance needed to improve to earn additional compensation, and ii) a monitoring-pressure condition, where participants were told that their performance was being videotaped and would be watched by researchers ^25^. These researchers reported that while the outcome-pressure and control conditions did not differ in their performance during learning, the monitoring-pressure condition produced a significant detriment in performance. Thus, if participants engage in greater performance monitoring during the intentional learning condition, they might exhibit a similar deficit in performance.

In this study, we observed that the effect of feedback differed over the course of the learning period for participants in the unintentional learning condition. The punishment group showed greater sequence knowledge expression compared to the reward and control groups in the first and last learning blocks, but not during the middle blocks of the learning period. The time course of sequence knowledge expression may be due to the interleaving of fixed- and random-sequence blocks during the learning period. In the interleaved design, participants may undergo forgetting or unlearning of the sequence during the random-sequence blocks ^29-31^. In this study, training with punishment might enhance early learning during the first sequence block, but also quicken adaptive forgetting during the subsequent random block, as instances of negative reinforcement may serve as an implicit signal to shift strategies or explore new behaviors (in this case, to “forget” any previously learned sequence knowledge). In contrast, participants training with reward may be less sensitive to slight changes in reinforcement frequency and be less likely to shift rules or undergo forgetting, leading to the development of sequence knowledge over a longer period of exposure. This pattern of behavior is consistent with a win-stay, lose shift strategy ^32,33^. Notably, it has been shown that reward is not effective in shaping motor behaviours unless participants are aware of the manipulation being rewarded or if reward is too abundant ^34,35^, probably due to participants not engaging in exploratory behaviors. Future work may also consider the psychological implications of reward and punishment given during learning, and how this interacts with behavioral variability and cognitive strategies.

The design implemented here precludes modelling the cognitive strategies, such as the win-stay lose-switch versus reinforcement learning strategies ^33^. However, based on the role of dopamine in perseveration, i.e., the use of reinforcement-learning strategy, we are inclined to speculate that punishment may promote participants to update their sequence-item knowledge more rapidly (effectively facilitating a win-stay lose-switch strategy), while reward may facilitate greater use of a reinforcement-learning strategy and integrate their sequence-knowledge over a longer period of time ^36^.

#### Recruiting different neural resources

Besides cognitive load, one potential explanation for the lack of benefit of punishment to performance is that learning with punishment and explicit sequence learning utilize shared neural substrates, and therefore compete for these resources. While our current data cannot speak directly to the neural basis of this effect, our prior work may provide insight into this question^37^. Explicit sequence learning is known to recruit the hippocampus ^38^, and our previous fMRI experiment provides evidence that training with punishment also engages medial temporal lobe, while reward recruits the cortical motor network ^37^. It is possible that these two processes interact competitively at the level of medial temporal lobe, thereby reducing the expression of early sequence knowledge. This possibility may be interesting to explore in the future using patients with medial temporal lobectomy ^39^.

### Reward and punishment did not impact immediate retention

Reward did not benefit performance on the immediate retention probes in the intentional or the unintentional learning conditions, which effectively replicates the similar null result found in prior work by our group ^6^. One notable difference between the present study and our prior work as that we did not include a delayed retention test, and so it is possible that reward may have benefitted delayed retention or offline gains. However, in our previous work^6^ wherein we did measure performance after delay, we found no effect of feedback on retention. Given the widely observed benefit of reward to visuomotor adaptation memory retention, it will be important to understand whether any specific conditions reliably foster reward-related memory benefits in the context of sequence learning ^4,19-21^.

### Intention and awareness

One might ask whether it is possible to between distinguish intention and awareness. In the context of movement, fMRI evidence suggests that intention and awareness are separable at least at the neural level ^40,41^. Additionally, our study demonstrates that intention and awareness are dissociable behaviorally. Based on the PDP several participants in the Unintentional learning condition demonstrated inclusion greater than chance-level, meaning that they had become aware of the sequence. On the other hand, multiple participants in the Intentional learning condition performed at or below chance level on the PDP, suggesting that they had no awareness of the sequence. This pattern of results demonstrates that awareness is dissociated from intention, although intention may bias implicit or explicit knowledge formation.

Historically, in a typical SRTT study of explicit and implicit knowledge, intention may be conflated with awareness ^8,9,42-49^. Specifically, participants in an “explicit learning condition” will be made aware of the sequence and might be directed to learn the sequence, while participants in the “implicit learning condition” will not be made aware of the sequence and simply be told to press buttons. Despite the condition being assigned prior to learning (i.e. implicit or explicit conditions), awareness is assessed post-hoc: participants are classified as having acquired knowledge implicitly or explicitly on the basis of the awareness test ^5,8-10,43-46,49,50^. At the end of learning, the participants in the explicit condition have *intended to learn* and may (or may not) end up being *aware of the sequence* they have been exposed to. On the other hand, the participants in the implicit condition learn *unintentionally*; however, it is possible that they will end up *being aware of the sequence*. Because participants are generally excluded from the experiment if they fail to show the expected type of sequence knowledge (explicit or implicit knowledge, e.g. Wachter et al. 2009 ^5^), these studies end up preserving the intended grouping. However, we suggest that the difference in instruction also may lead to an unstated difference between the conditions: namely, that they had differing intentions during training.

### Considerations

Our conclusion that intention impacts the influence of punishment on early sequence learning is drawn from the direct comparison of the performance during the first training block (i.e. S1 and R1) in the Intentional and Unintentional learning conditions. This comparison was motivated by our results ^6^, which revealed that both punishment and reward led to better sequence knowledge expression early in learning. This comparison was therefore hypothesized a *priori* - we note that the omnibus ANOVA that included all training blocks (i.e. S1-S8 and R1-R8) did not show a significant interaction between Intention × Feedback × Block (p = 0.055). However, our study was appropriately powered to detect the difference between the groups observed in the initial block, where we hypothesized the difference between the feedback conditions would be observed. Because we detected the benefit of punishment as we expected, and effectively replicated our prior results, we are confident in our conclusion that punishment benefits early sequence knowledge expression when learning unintentionally.

## Conclusions

One question that the current study does not address is the extent to which the intentional learning condition should be considered a separate task to unintentional learning. Implicit and explicit sequence learning are often considered together in the same studies and treated as equivalent conditions ^8,9,48,51^, and so there is precedent in the literature to consider these types of manipulations as equitable variations of the same task. This assumption is primarily based on task demands; the two tasks are functionally equivalent, in the sense that the equivalent stimulus elicits the same response in both cases, and, importantly, the contingency to receive the reward or punishment is equivalent between the two tasks. Given these considerations although these variations of the task undoubtedly engage two different cognitive processes, the intentional and unintentional learning conditions may be comparable tasks.

In conclusion, in the present study we demonstrated that punishment may enhance early sequence knowledge expression (on the SRTT) specifically when participants learn unintentionally. As is the case with visuomotor adaptation, the effect of valenced feedback in sequence learning may also depend on explicit processes. Future work should consider other factors that might influence the impact of reward, including the participant’s psychological state prior to the experiment, their personality, and the time of day.

## Methods

### Participants

72 participants were recruited for this study. 36 participants were assigned to each experiment (Experiment 1: 18 female, age = 24.4±3.1 [mean ± standard deviation]; Experiment 2: 17 female, age = 23.3±2.4; Figure 1a), such that 12 participants were assigned to each feedback group. We chose 12 participants in each group based on a power analysis of the difference in feedback conditions during early learning observed in our previous work ^6^. There was no difference between the mean age of the participants in Experiments 1 and 2 (t_(70)_ = 1.78, p = 0.0784). All participants were right-handed, free from neurological or psychiatric disorders, and had normal or corrected-to-normal vision. All participants gave informed consent and the study was performed with approval from the Oxford Central University Research Ethics Committee (R44415/RE001), and the study was run in accordance with the Declaration of Helsinki. Participants were compensated for their time in local currency (GBP, £).

### Experimental Outline

Participants trained on the serial reaction time task (SRTT) over 30 blocks. Each block contained 48 trials (Figure 1b). During some blocks (“fixed-sequence blocks” [S]) the sequence of stimuli would appear according to a repeating pattern (described below for each task). During other periods, stimuli appeared in a pseudo-randomly determined order (“random-sequence blocks” [R]). During the training period, fixed- and random-sequence blocks were presented in an interleaved fashion ^52^, to make gaining explicit knowledge more challenging, and to allow us to assess the development of sequence knowledge (compared to general skill) continuously across the learning period.

In both experiments, the task began with three random-sequence blocks without feedback (“familiarisation blocks”, F1, F2, F3) to familiarise participants to the task and to establish their baseline level of performance. The subsequent training period also began with a random block (R0), after which blocks alternated between random and sequence blocks (S1-R8, total number of blocks = 16). The training period concluded with a random block. The difference in performance between an S-block and the subsequent R-block (e.g., S1 – R1, S2 – R2, and so on) was used to index sequence knowledge during the training period ^8^. Immediately following completion of the training period, participants were tested for sequence memory without feedback during the probe period. The probe period consisted of 9 blocks with either a fixed sequence (Probe fixed-sequence [PS]) or a random sequence (Probe random-sequence [PR]) without feedback in the following order: PR – PS – PR – PS – PS – PS – PS – PR – PS – PR. Sequence knowledge was indexed by comparing the initial probe sequence block (PS1) to the subsequent random block (PR2). The four fixed sequence blocks after the first sequence knowledge test (PSt 1-3 and PS2) were included to determine whether subjects trained with punishment relearn at a faster rate, which has been suggested by prior research using visuomotor adaptation tasks ^19^. The probe period concluded with a second sequence knowledge test to determine whether the groups differed in the sequence knowledge after the prolonged sequence period.

To test the impact of reward and punishment on skill learning, participants were randomized into one of three Feedback groups: reward, punishment, or uninformative (control). During the feedback period, reward, punishment, or control feedback was provided based on the participant’s ongoing performance.

### Serial reaction time task

The version of the SRTT used here adds feedback to the traditional implementation. At the beginning of each block participants were presented with four “O”s, arranged along a horizontal line at the centre of the screen. These stimuli were presented in white on a grey background. A trial began when one of the “O”s changed to an “X”. Participants were instructed to respond as quickly and accurately as possible, using the corresponding button, on a four-button response device held in their right hand. The “X” disappeared once the subject made a response, followed by a 200 ms fixed inter-trial interval, during which time the four “O”s were displayed.

A block consisted of 48 trials. During fixed-sequence blocks, the stimuli appeared according to a fixed 12-item sequence repeated 4 times. The four sequences used in this study were taken from our previous study^6^: *1) 2,4,2,1,3,4,1,2,3,1,4,3; 2) 3,4,3,1,2,4,1,3,2,1,4,2; 3) 3,4,2,3,1,2,1,4,3,2,4,1; 4) 3,4,1,2,4,3,1,4,2,1,3,2*. Participants were randomly assigned to one of the sequences. Each fixed block began at the same position in the sequence. In the random blocks, the stimuli appeared according to a pseudo-randomly generated sequence, without repeats on back-to-back trials, so, for example, participants would never see the triplet 1-1-2. Each block lasted roughly one minute, and the entire experimental session lasted approximately 35 minutes.

Between each block, participants were presented with the phrase “Nice job, take a breather.” After five seconds, a black fixation-cross appeared on the screen for 25 seconds. Five seconds before the next block began, the cross turned blue to cue the participants that the block was about to start.

The first button press made after stimulus presentation was considered the participant’s response. Only correct trials were considered for analysis of RTs. Trials with RTs less than 150 ms and greater than 800 ms were excluded from the analysis. The bounds of this exclusion criteria are consistent with prior work from our lab ^6,53^.

### Intentionality manipulation

In Experiment 1, after the familiarisation period, participants were given the following instructions, “During certain periods of time, you may have the feeling that the stimulus is following a fixed sequence. Please ignore the sequence and continue to perform as fast and accurately as possible.”

In Experiment 2, participants received the following instructions, “During certain periods of time, you may have the feeling that the stimulus is following a fixed sequence. Please do your best to learn the sequence, while also performing as fast and accurately as possible.”

Participants were not given any further instructions about the nature of the task. Thus, we refer to the participants in Experiment 1 as “unintentional learners” and those in Experiment 2 as “intentional learners.”

### Valenced Feedback

All participants were paid a base remuneration of £15 for participating in the study. At the conclusion of the familiarisation period, participants were told they could earn more money based on their performance. Participants were randomly assigned into the reward, punishment, or control feedback groups. The presence of reward or the absence of punishment was based on participant’s performance. In both versions of the SRTT, an initial criterion RT was defined based on the participant’s median performance during the final familiarization block. As participants progressed through training, this criterion was re-evaluated after each block, to encourage continuous improvement. In the reward group, the feedback indicated that the participant’s performance was improving. In the punishment group, the feedback indicated they were getting worse. The control group received feedback on 50% of trials that were randomly determined. Participants in the control group were told that the feedback was not meaningful. We considered the reward and punishment control groups together in the analyses, as is typical in these studies ^5,11^.

Performance was defined as the accuracy (correct or incorrect) and RT of a given trial. Feedback was given on a trial-by-trial basis and was indicated to the participant by the white frame around the stimulus changing to either green (reward) or red (punishment). In the reward group, the participants were given feedback if their response was accurate and their RT was faster than their criterion RT, which indicated that they earned money (£0.02 from a starting point of £0) on that trial. In the punishment group, participants were given feedback if they were incorrect, or their RT was slower than their criterion, which indicated that they lost money (£0.02 deducted from a starting point of £20) on that trial. Participants in the control group saw red or green colour changes on 50% of trials that were randomly determined (and therefore unrelated to their performance).

The initial criterion RT was calculated as median performance in the first familiarization block (F1). After each block, the median + standard deviation of performance was calculated and compared with the criterion. If this test criterion was faster than the previous criterion, the criterion was updated. Only correct responses were considered when establishing the criterion RT.

To ensure they were adequately motivated, participants in the control group were told that they would be paid based on their speed and accuracy. Importantly, to control for the motivational differences between gain and loss, participants were not told the precise value of a given trial. This allowed us to assess the hedonic value of the feedback, rather than the level on a perceived-value function. For the reward and punishment groups, the current earning total was displayed (e.g., “You have earned £5.00”) between the blocks. For the control group the phrase, “You have earned money.” was presented.

### Conscious recall

In both experiments to assess conscious recall, we used both verbal report and the process dissociation procedure ^10^. For the verbal report, participants reported as much of the sequence as they knew. The total number of correct sequential items verbally reported was compared using a one-way ANOVA with Feedback type (Reward/Control/Punishment) as a factor for each experiment separately. We also confirmed that the Intentional instructions fostered more explicit knowledge than the Unintentional instructions using a two-way ANOVA with Feedback type (Reward/Control/Punishment) and Intentionality (Intentional/Unintentional) as factors.

For the process dissociation procedure, after completion of the probe blocks, participants performed 3 sets of 100 button presses. First, we asked participants to generate 100 button presses randomly with no repeats. At this point, the presence of the sequence was confirmed, and participants were then asked to input 100 button-presses while trying to repeat as much of the sequence as they could recall. Participants were specifically instructed that if they thought they knew any of the sequence, or several chunks of the sequence, that they should input those pieces as many times as they could. Participants were then asked to generate 100 button presses where they tried to exclude the sequence as much as possible. The ‘random generation’ test for the process dissociation procedure tested the hypothesis that reward may prime participants to perform the sequence more readily even during random button pressing. In addition, the exclusion condition was used to assess conscious ‘control’ over excluding the sequence, as has been implemented previously in the literature ^7,10,54^. The area under the curve for each number of sequential items (triplets, quadruplets, quintuplets through dodecs) included in the 100 button presses generated by the subject was compared to chance, which was estimated via 10,000 Monte-Carlo simulations of 100 non-repeating items with the probability of occurrence uniformly distributed across the 4 buttons. The area under the curve for each group was compared using a one-way ANOVA with Feedback type (Reward/Control/Punishment) as a factor.

### Statistical analysis

Data analysis was conducted using JASP (Version 0.8.4, for Mac). To ensure there were no initial differences between the groups, the mean RT during the familiarization blocks were compared using a one-way ANOVA with Feedback type (Reward/Punishment/Control) as a factor. To evaluate the impact of reward and punishment during the training period, RTs were compared across the groups using a repeated measures ANOVA with Feedback type (Reward/Punishment/Control), Sequence type (Fixed-/Random-sequence), and Block number (1-8) as factors. For the probe period, the first and second sequence knowledge tests were compared using a repeated measures ANOVA with Feedback type (Reward/Punishment/Control), Sequence type (Fixed-/Random-sequence), and Test (Test 1/Test 2) as factors. To test for differences in relearning, the four consecutive sequence blocks during the probe period were compared using a repeated measures ANOVA model with Feedback type (Reward/Punishment/Control) and block (PSt1-3 & PS2) as factors.

To formally test whether the learning rate differed between the unintentional and intentional experiments, RTs were compared across the groups using a repeated measures ANOVA with Experiment (Intentional/Unintentional), Feedback type (Reward/Punishment/Control), Sequence type (Fixed-/Random-sequence), and Block number (1-8) as factors. Because we were specifically interested in early learning, we also considered the effect of feedback specifically during early learning (Block 1) with a repeated measures ANOVA with Experiment (Intentional/Unintentional), Feedback type (Reward/Punishment/Control), Sequence type (Fixed-/Random-sequence) as factors.

Post-hoc tests were conducted using t-tests, with multiple comparisons corrected using the Bonferroni-Holm method where appropriate.

## Acknowledgements

AS and CIB are funded by the NIMH internal research program (ZIA-MH002893). CJS holds a Sir Henry Dale Fellowship, funded by the Wellcome Trust and the Royal Society (102584/Z/13/Z). The Wellcome Centre for Integrative Neuroimaging is supported by core funding from the Wellcome Trust (203139/Z/16/Z). AS would like to thank the International Biomedical Research Alliance for their support.

## Conflict of interest

The authors declare no conflicts of interest.

## Author contributions

AS, CB, and CJS designed the study. AS collected and analysed the data. AS, CB, and CJS wrote the paper.

## Supplemental information

**Table 1.**
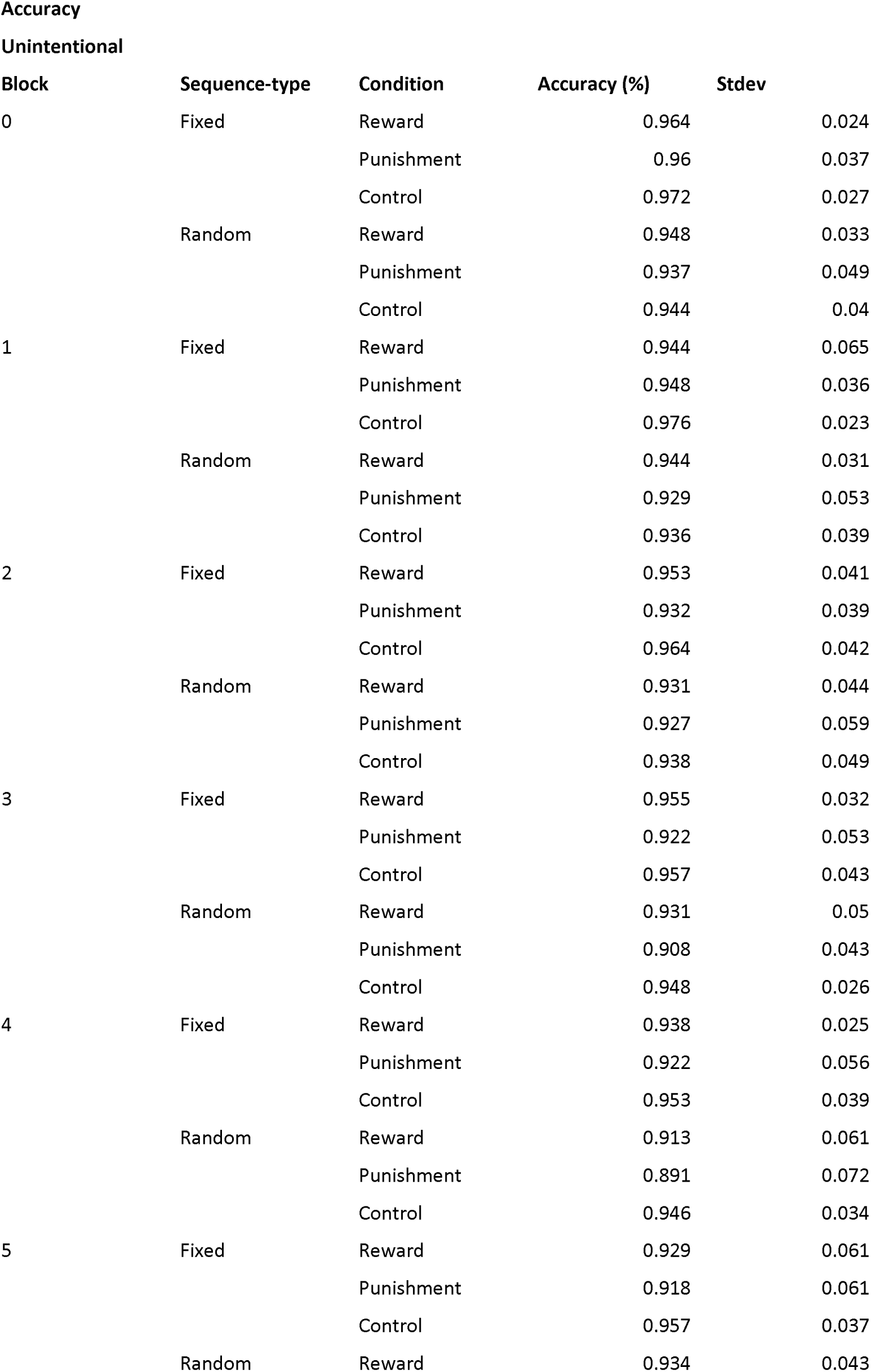
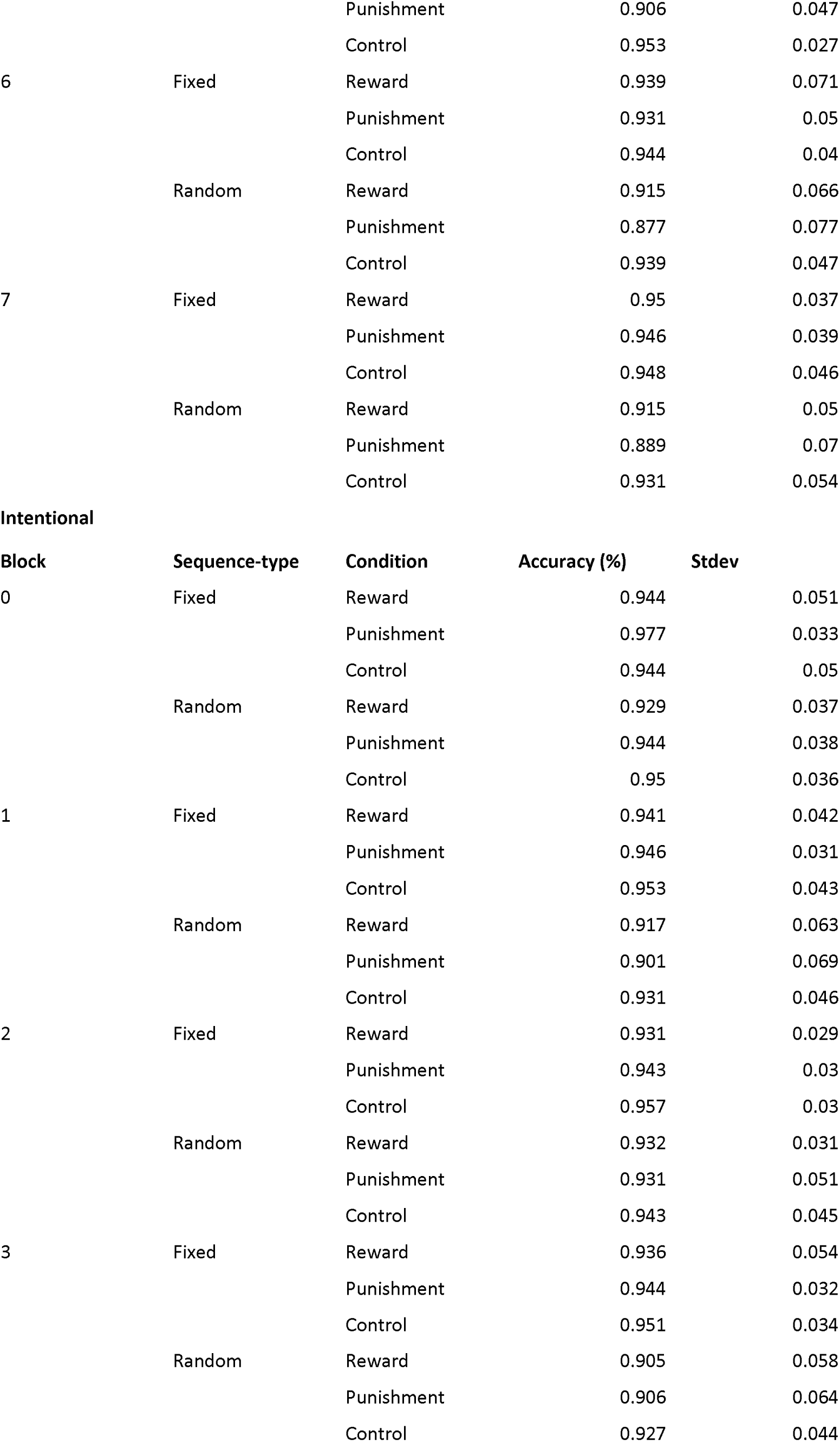
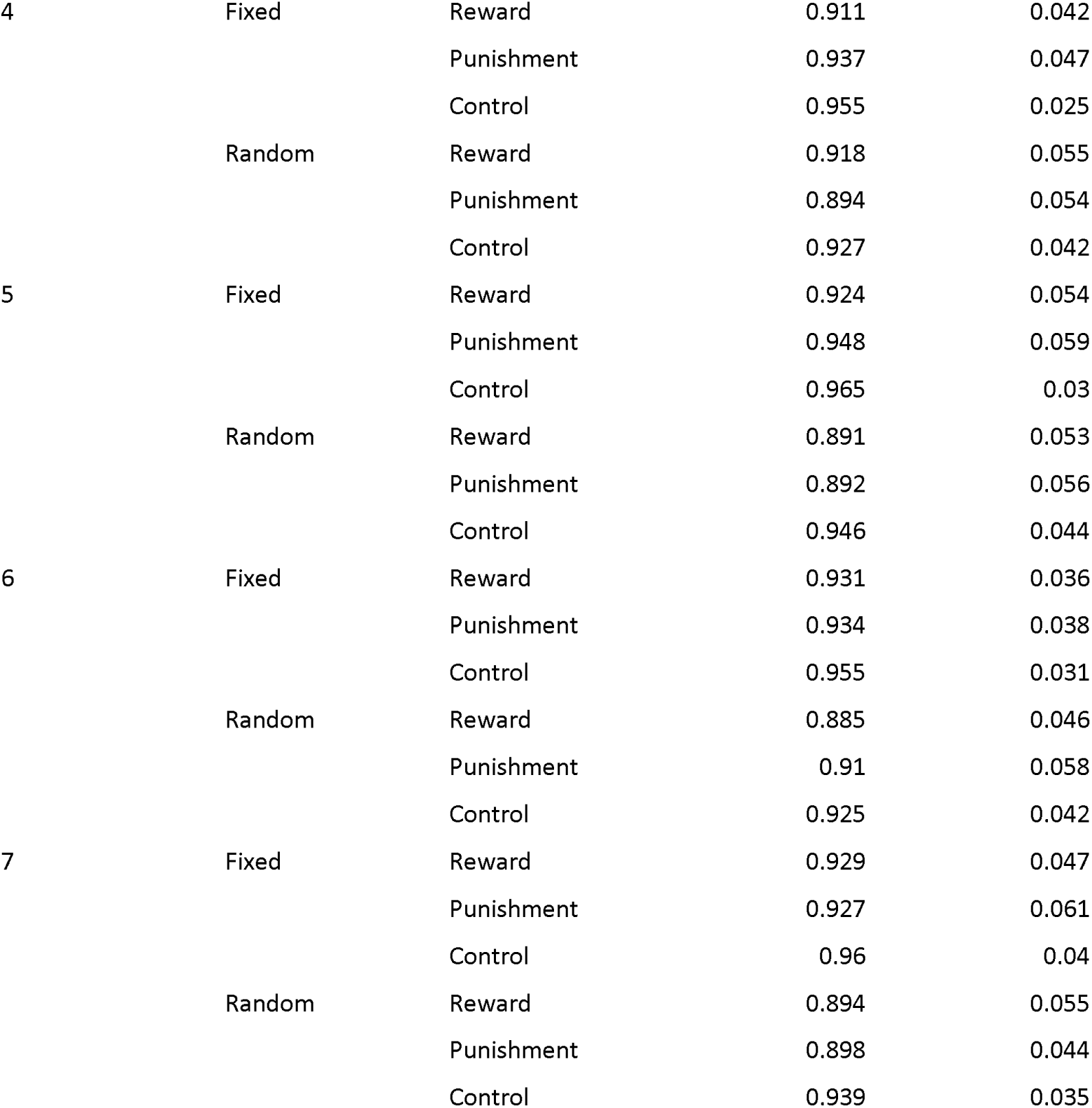
Accuracy data for the intentional and unintentional experiments.

## Notes

#### Summary of Updates

Added discussion points re: awareness and statistical analyses

